# Hospitalizations associated with respiratory syncytial virus (RSV) and influenza in children, including children having a diagnosis of asthma

**DOI:** 10.1101/161067

**Authors:** E. Goldstein, L. Finelli, A. O’Halloran, P. Liu, Z. Karaca, C. Steiner, C. Viboud, M. Lipsitch

## Abstract

**Background:** There is uncertainty about the burden of hospitalization associated with RSV and influenza in children, including those with underlying medical conditions.

**Methods:** We applied previously developed methodology (Goldstein et al., Epidemiology 2012) to HealthCare Cost and Utilization Project (HCUP) hospitalization data and additional data related to asthma diagnosis/previous history in hospitalized children to estimate RSV and influenza-associated hospitalization rates in different subpopulations of US children between 2003-2010.

**Results:** The estimated average annual rates (per 100,000 children) of RSV-associated hospitalization with a respiratory cause (ICD-9 codes 460-519) present anywhere in the discharge diagnosis were 2381 (95% CI(2252,2515)) in age <1y; 710.6(609.1,809.2) (age 1y); 395(327.7,462.4) (age 2y); 211.3(154.6,266.8) (age 3y); 111.1(62.4,160.1) (age 4y); 72.3(29.3,116.4) (ages 5-6y); 35.6(9.9,62.2) (ages 7-11y); and 39(17.5,60.6) (ages 12-17y).

The corresponding rates of influenza-associated hospitalization were lower, ranging from 181(142.5,220.3) in age <1y to 17.9(11.7,24.2) in ages 12-17y. The relative risks for RSV-related hospitalization associated with a prior diagnosis of asthma in age groups under 5y ranged between 3.1(2.1,4.7) (age <1y) to 6.7(4.2,11.8) (age 2y); the corresponding risks for influenza-related hospitalization ranged from 2.8(2.1,4) (age <1y) to 4.9(3.8,6.4) (age 3y).

**Conclusions:** RSV-associated hospitalization rates in young children are high and decline rapidly with age. Young children with an asthma diagnosis should be target groups for RSV and influenza-related mitigation efforts, possibly including RSV prophylaxis for the youngest children.

## Introduction

Robust estimation of severity is essential for planning prevention programs. However, there is uncertainty about the rates of hospitalization associated with RSV and influenza in different age groups of children, as well as in children with underlying medical conditions such as asthma. Studies of hospitalizations associated with respiratory virus infections in select communities (e.g. [1–5]) have documented high rates of RSV-associated hospitalization in full- and pre-term infants ([1,2]) and young children [1], as well as high rates of influenza-associated hospitalization in children during certain influenza seasons (e.g. [3] vs. [4]). Moreover, these studies reveal a high frequency of underlying medical conditions (particularly asthma) among children hospitalized with RSV and influenza virus infections ([1,5-7]). However, community-based studies can suffer from limitations such as under-detection of the presence of viral infections, and suboptimal sample sizes. Estimates of hospitalization rates associated with respiratory viruses can also be derived from studies using hospital discharge data. Limitations of discharge diagnosis studies may include both under-ascertainment and misclassification of specific respiratory viral etiologies. Strategies for correction of missing or misclassified diagnoses have been developed to produce burden estimates [8]. Statistical inference based on regression models for time series of severe outcomes such as deaths or hospitalizations is a common approach for estimating the contribution of respiratory viruses to the burden of these outcomes [9–14]. However, various assumptions underlying such statistical models may be uncertain [15,16], while the model estimates and the goodness-of fit are sensitive to those assumptions (e.g. Supporting Information for [13,14]). Reducing uncertainty in both the assumptions and burden estimates for respiratory virus infections, and increasing the specificity of estimates by narrow age group and by underlying condition could support the development and targeting of prevention and treatment. For example, a variety of RSV vaccine candidates for different populations are currently in different stages of development [17–19], but target groups for vaccination beyond infancy have not been well described.

Prevalence of asthma diagnosis in young children increases with age ([20]; Table 3). Diagnosis of asthma in infants and young children is challenging [21], with different causal mechanisms involved in the pathogenesis of asthma-like symptoms during childhood [22]. Nonetheless, several studies [1,5-7,23,24] have suggested that asthma diagnosis in children serves as a marker for the risk of RSV- and influenza-associated hospitalization. However, the age-specific magnitudes of those risks for the subpopulations of US children with an asthma diagnosis compared to the general population of children are not well established, particularly for RSV-associated hospitalizations. Quantification of those risks could therefore help inform RSV and influenza-related prevention and treatment strategies for those children.

In our earlier work [13,14], we introduced a new method for estimating the burden of severe outcomes associated with influenza and RSV, designed to address several limitations of some of the previously employed inference models. Important features of that approach include: the use of RSV and influenza (sub)type (A/H3N2, A/H1N1, and B) incidence proxies that are expected to be linearly related to the population incidence of those viruses; a flexible model for the baseline of severe outcomes not associated with influenza and RSV; and a bootstrap method for inferring the confidence bounds on the estimates of the rates of influenza and RSV-associated severe outcomes that accounts for auto-correlation in the time series for the noise.

Here, we apply the inference method in [13,14] to data on hospitalizations from US states that reported to the HealthCare Cost and Utilization Project (HCUP) for the 2003-04 through the 2009-10 seasons to better estimate the rates of influenza and RSV-associated hospitalization in US children. We estimate these rates for several categories of ICD-9 coded hospital discharge diagnoses. Our main goals are to provide detailed age-specific estimates (by year of age for the youngest children) of the incidence of influenza and RSV-associated hospitalization in the general population of children, as well as in the population of children having a diagnosis of asthma.

## Methods

### Hospitalization Data

We used weekly data on hospitalizations with different diagnoses in different age groups of children (<1y,1y,2y,3y,4y,5-6y,7-11y,12-17y) from the State Inpatient Databases of the Healthcare Cost and Utilization Project (HCUP), maintained by the Agency for Healthcare Research and Quality (AHRQ) through an active collaboration [25]. Thirty-five states reported to HCUP for the 2003-04 through the 2009-10 seasons. For our analyses, we included 24 states for which data on hospitalizations in infants with RSV in the principal diagnosis exhibited consistency throughout the study period (Supporting Information). Those 24 states, which constituted approximately 65.8% of the US population during the study period, are Arkansas, California, Colorado, Connecticut, Georgia, Hawaii, Iowa, Illinois, Indiana, Maryland, Minnesota, North Carolina, Nebraska, New Jersey, Nevada, New York, Ohio, Oregon, Tennessee, Texas, Virginia, Vermont, Washington, and Wisconsin.

Aggregate hospitalization data were used in our analyses, with no informed consent from participants sought.

### Statistical Inference Framework

Our study period is between calendar week 27 of 2003, and calendar week 26 of 2010.

As the incidence rates of influenza and RSV infection are difficult to estimate directly, we use *proxies* for the incidence of RSV and the major influenza (sub)types (A/H3N2, A/H1N1, and influenza B) that are expected to be proportional to the population incidence of those viruses [13–15]. Those proxies are derived from the available data as described below.

For the influenza incidence proxies, we utilized the US Centers for Disease Control and Prevention (CDC) influenza surveillance data [26]. We multiplied the weekly state-specific percent of medical consultations in the CDC Outpatient Illness Surveillance Network (ILINet) that were for influenza-like illness (ILI) by the state-specific percent of respiratory specimens in the US Virologic Surveillance laboratories that tested positive for each of the major influenza (sub)types (A/H3N2, A/H1N1, and influenza B) to define state-specific weekly proxies for the incidence of each major influenza (sub)type:

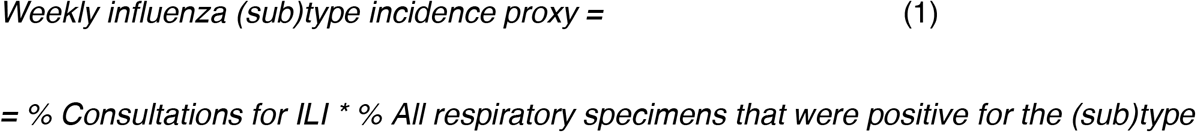

Incidence proxy for an influenza (sub)type for the collection of twenty-four US states used in our analyses (Data subsection) is defined as the sum of state-specific incidence proxies for the (sub)type weighted by the weekly state populations. Weekly state populations during our study period were estimated through linear interpolation (in time) for the yearly, July 1st state population estimates [27]. Finally, the relation between an influenza incidence proxy for a given influenza (sub)type and the rate of associated hospitalization in a given age group may change over time due to a variety of factors, particularly the circulation of novel influenza strains (see [13], as well as Supporting Information). To accommodate those changes, we split the incidence proxies for both influenza A/H3N2 and A/H1N1 into several time periods – see Supporting Information for full details.

The weekly RSV incidence proxy used in this study was the cumulative rate of hospitalization among infants with RSV (ICD-9 codes 466.11, 480.1, or 079.6 present in the principal discharge diagnosis) during that week for the collection of states included in our analyses. We use hospitalization rates in infants because there is an apparent upward trend in annual rates of hospitalization with the principal diagnosis of RSV in all age groups above two years of age in the US, likely due to changes in testing/diagnostic practices, but no apparent trend in the rates of hospitalization with the principal diagnosis of RSV in infants. We note that the use of the above RSV incidence proxy is premised on the assumption that there is year-to-year consistency in the relation between RSV infection rates in different age groups of children. Such temporal consistency may not hold for the relation between RSV infection rate in children vs. adults, and the use of the above incidence proxy for estimating hospitalization rates in adults may be questionable. We refer to Section S2 of Supporting Information for further details on the RSV incidence proxy.

The outcomes we evaluated are rates of hospitalization for six different diagnosis categories (Table S1 in the Supporting Information) by age group in children. Those categories are 1) respiratory cause excluding asthma (ICD-9 codes 460-519; excluding 493) in the principal discharge diagnosis; 2) respiratory cause (ICD-9 codes 460-519) present anywhere in the discharge diagnoses (principal or secondary) excluding asthma (ICD-9 code 493) in the principal diagnosis; 3) respiratory cause (ICD-9 codes 460-519) present anywhere in the discharge diagnosis; 4) pneumonia and influenza (ICD-9 codes 480-488) in the principal discharge diagnosis; 5) asthma (ICD-9 code 493) as a secondary (non-principal) discharge diagnosis; 6) asthma (ICD-9 code 493) present anywhere in the discharge diagnosis. The first three categories are used to evaluate the burden of RSV and influenza-related respiratory hospitalizations, from the more restricted (category 1) to the most inclusive category of hospitalizations (category 3). Category 4 represents an important subcategory of respiratory hospitalizations. Categories 5 and 6 are subsequently used to estimate the risks for RSV and influenza-related hospitalization associated with having a prior diagnosis of asthma.

Hospitalization Rate Estimation: For each choice of an age group and diagnosis category 1-6, weekly hospitalization rates in that age group with that diagnosis category during the study period (*outcome*) are regressed linearly against the following covariates: incidence proxies for the different influenza (sub)types and RSV, as well as temporal trend (modeled by a low degree polynomial in time) and a seasonal baseline with annual periodicity (modeled by cubic splines in the calendar week with knots at every fourth week, periodic in calendar year) [13,14]. If *O*(*t*) is the weekly rate for the outcome on week *t*, and *V_i_* (*t*) are the incidence proxies for the various viruses, namely the different influenza (sub)types (split into time periods as described in the Supporting Information) and RSV, the model structure is

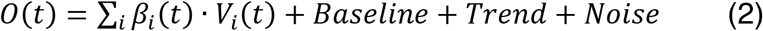

To account for the autocorrelation structure in the noise, a bootstrap method devised in [13] is used to estimate the confidence bounds for the model’s estimates. Estimates derived in earlier work [14] suggest that several alternative inference methods (such as maximal likelihood estimation under the AR(1) assumption for the noise structure) yield very similar results.

Finally, we note that rates of influenza-associated hospitalization for categories 3 and 6 are estimated differently for reasons described in Section S3 of the Supporting Information.

### Risks for RSV and influenza hospitalization associated with a prior diagnosis of asthma

Data on the prevalence of asthma diagnosis by year of age between 2003-2009 were provided to us by Dr. L. Akinbami of the US CDC (see also [20]).

To estimate the relative risk for influenza hospitalization in a given age group associated with a prior diagnosis of asthma we estimate

i. Age-specific proportion *H*(*age*) of cases of influenza-related hospitalizations in the Influenza Hospitalization Surveillance Network FluSurv-NET surveillance database [28,7] between 2003-04 through 2009-10 seasons that have a previous history of asthma, including reactive airway disease.
ii. Age-specific population prevalence *D*(*age*) of asthma diagnosis between 2003-2009 [20].

The relative risk (risk ratio) *RR_Rflu_* (*age*) for influenza hospitalization in a given age group associated with a prior diagnosis of asthma is then estimated as:

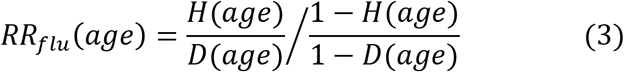

Estimation of the (age-specific) relative risk for RSV hospitalization associated with having a prior diagnosis of asthma is described in Section S4 of the Supporting Information.

## Results

Figure 1 shows the model fits for the category of hospitalizations with a respiratory cause present anywhere in the discharge diagnosis in children aged <4y, as well as the contribution of both RSV and influenza to the rates of these hospitalizations. Figures 2 and 3 show the corresponding information for the categories of hospitalizations with pneumonia and influenza in the principal diagnosis, and asthma as a secondary (non-principal) diagnosis. Further model fits are presented in the Supporting Information. All those figures show good, temporally consistent model fits (R-squared statistic above 0.985 for all models in Figures 1-3, Supporting Information), as well as the major contribution of RSV to the corresponding hospitalization burden.

**Figure 1:**
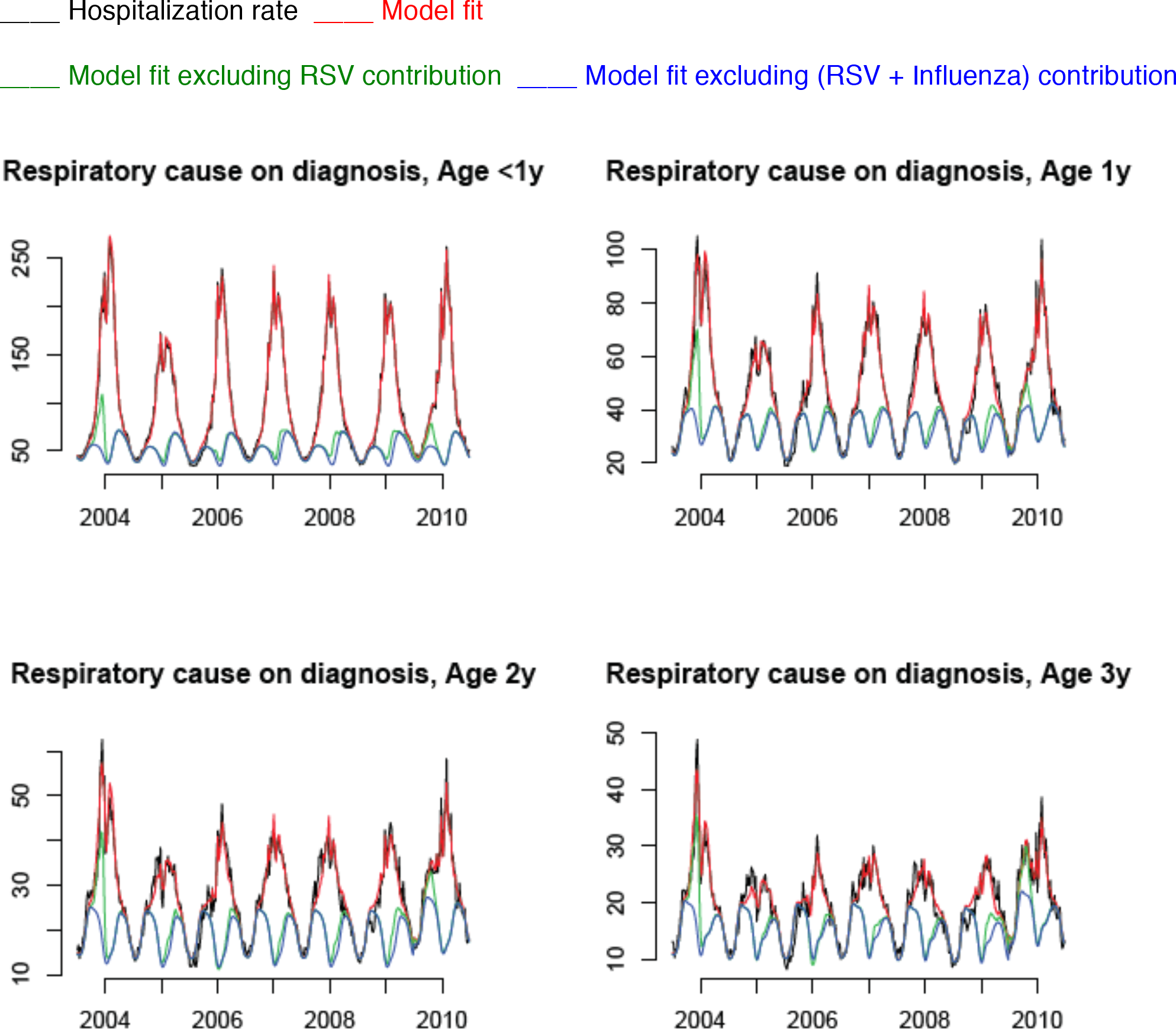
Weekly hospitalization rates (per 100,000) with a respiratory cause present anywhere in the discharge diagnosis for children aged <4y, 2003-04 through the 2009-10 seasons (black), model fits (red), and contributions of RSV (red curve minus green curve) and influenza (green curve minus blue curve).

**Figure 2:**
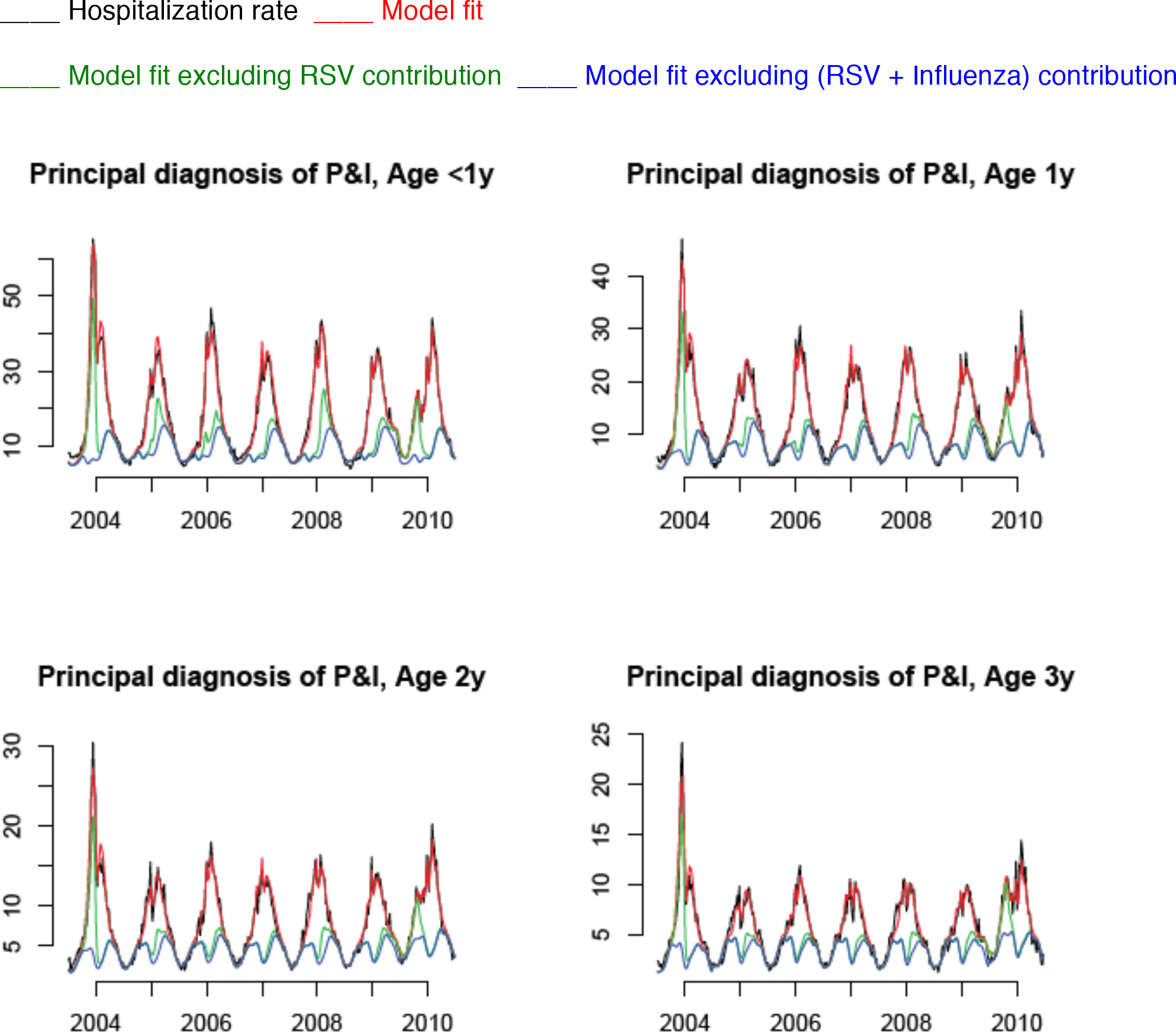
Weekly hospitalization rates (per 100,000) with pneumonia and influenza in the principal discharge diagnosis for children aged <4y, 2003-04 through the 2009-10 seasons (black), model fits (red), and contributions of RSV (red curve minus green curve) and influenza (green curve minus blue curve).

**Figure 3:**
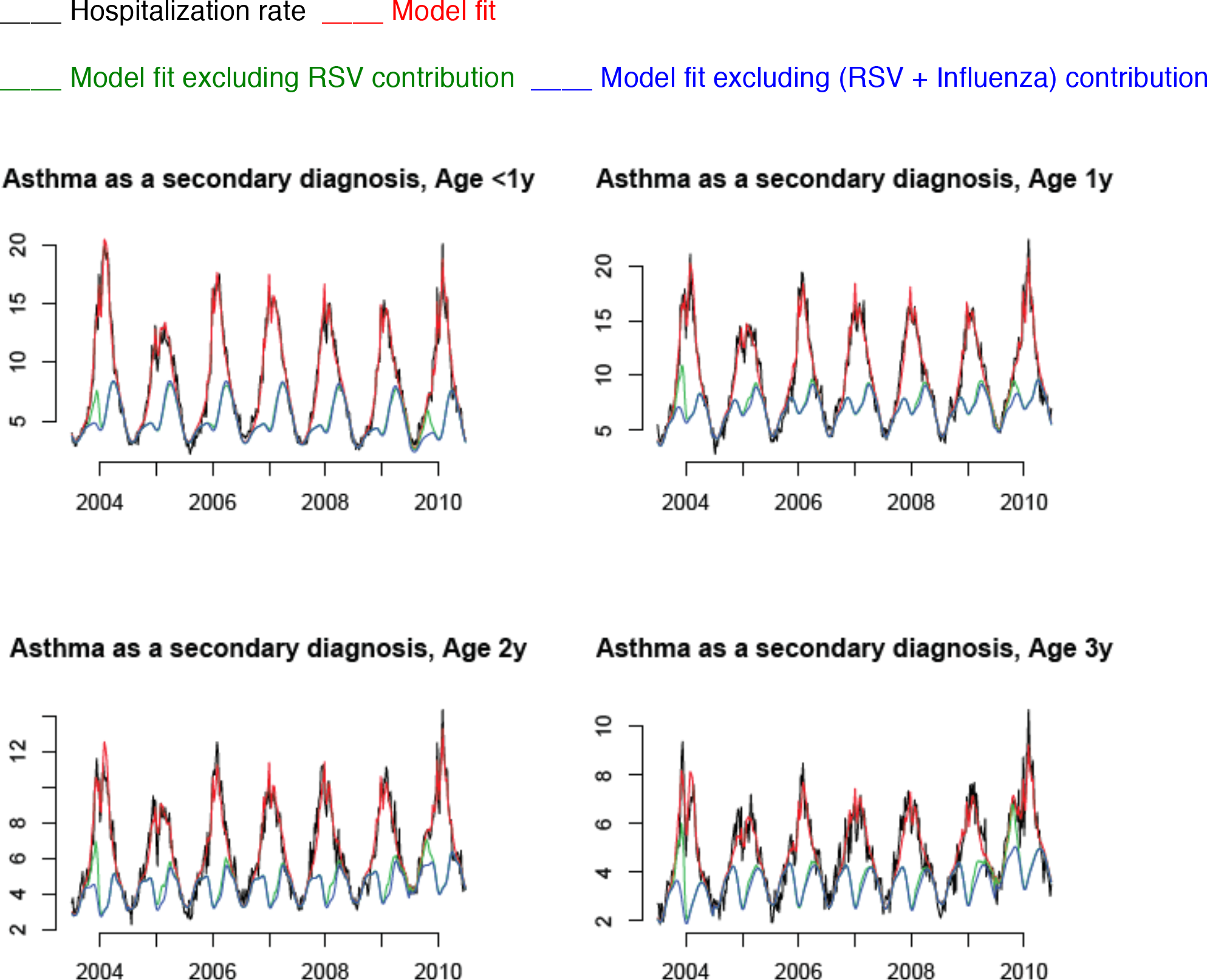
Weekly hospitalization rates (per 100,000) with asthma as a secondary (non-principal) discharge diagnosis for children aged <4y, 2003-04 through the 2009-10 seasons (black), model fits (red), and contributions of RSV (red curve minus green curve) and influenza (green curve minus blue curve).

Tables 1 and 2 present the estimates of the average annual rates of hospitalization associated with influenza and RSV for the six categories of discharge diagnoses (Methods) in select age subgroups of children. For all age groups and diagnosis types studied (except P&I hospitalizations in children aged 7-17y), the estimates of the rates of RSV-associated hospitalization are higher than the estimates of the rates of influenza-associated hospitalization. The estimated rates of RSV-associated hospitalization with a respiratory cause present anywhere in the discharge diagnosis are highest in children aged <1y (Table 1), and decline rapidly with age. For the more restricted categories of respiratory causes present anywhere in the discharge diagnosis (excluding asthma in the principal diagnosis), and respiratory causes excluding asthma in the principal diagnosis, the rates of RSV-associated hospitalization were progressively lower (Table 1). A sizeable fraction of both RSV and influenza-associated hospitalizations with a respiratory cause present anywhere in the diagnosis in different age groups (except RSV hospitalizations in infants aged <1y) had P&I in the principal diagnosis (Table 2). Additionally, for all age groups of children except four year-olds, the majority of RSV and influenza-associated hospitalizations with asthma present anywhere in the discharge diagnosis have asthma as a secondary (non-principal) diagnosis (Table 2).

**Table 1.** Average annual rates of influenza and RSV-associated hospitalization with a respiratory cause (excluding asthma) in the principal discharge diagnosis (ICD-9 codes 460-519 excluding 493); a respiratory cause present anywhere in the discharge diagnosis (excluding asthma in the principal diagnosis); and respiratory cause present anywhere in the discharge diagnosis per 100,000 US children in different age groups, 2003-04 through the 2009-10 seasons.

| Age group | Hospitalizations with a respiratory cause (excluding asthma) in the principal diagnosis |  | Hospitalizations with a respiratory cause in either the principal or secondary diagnosis (excluding asthma in the principal diagnosis) |  | Hospitalizations with a respiratory cause present anywhere in the diagnosis |  |
| --- | --- | --- | --- | --- | --- | --- |
|  | Flu | RSV | Flu | RSV | Flu | RSV |
| <1y | 140.9<br>(114.1,168.4) | 2129<br>(2044,2214) | 179.3<br>(141.6,217.6) | 2347<br>(2224,2471) | 181<br>(142.5,220.3) | 2381<br>(2252,2515) |
| 1y | 66.3<br>(41.3,91.4) | 561.2<br>(483,639.4) | 80.7<br>(53.6,108.4) | 631.2<br>(543.9,716.9) | 85.6<br>(57.3,114.5) | 710.6<br>(609.1,809.2) |
| 2y | 41.2<br>(28.1,53.9) | 265<br>(223.2,307.3) | 57.4<br>(41.7,73.3) | 322.3<br>(272.8,372.2) | 62<br>(45.3,79) | 395<br>(327.7,462.4) |
| 3y | 32.1<br>(22.8,41.7) | 140.9<br>(111.2,171) | 44.5<br>(32.7,56.5) | 168.8<br>(131.4,207.2) | 47.7<br>(35.6,60.9) | 211.3<br>(154.6,266.8) |
| 4y | 26.5<br>(19.8,33.3) | 66.1<br>(44.3,88.1) | 36.8<br>(28.2,45.5) | 87.7<br>(59.3,116.2) | 41.3<br>(31.7,51.3) | 111.1<br>(62.4,160.1) |
| 5-6y | 27.6<br>(22.2,33) | 41.5<br>(23.6,59.2) | 36.8<br>(29.6,44) | 62.8<br>(39.3,86.3) | 40.2<br>(32.4,48.4) | 72.3<br>(29.3,116.4) |
| 7-11y | 14.7<br>(12.3,17.1) | 16.8<br>(9.2,24.5) | 20.4<br>(16.3,24.4) | 27.6<br>(14.1,41.1) | 23<br>(18.5,27.7) | 35.6<br>(9.9,62.2) |
| 12-17y | 9.9<br>(8.5,11.3) | 14<br>(9.3,18.8) | 15.9<br>(10.5,21.3) | 33.8<br>(15.2,52.6) | 17.9<br>(11.7,24.2) | 39<br>(17.5,60.6) |

**Table 2.** Average annual rates of influenza and RSV-associated hospitalization with P&I (ICD9 codes 480-488) in the principal discharge diagnosis, asthma as a secondary (non-principal) discharge diagnosis, and asthma present anywhere in the discharge diagnosis per 100,000 US children in different age groups, 2003-04 through the 2009-10 seasons.

| Age group | Pneumonia and Influenza in the principal diagnosis |  | Asthma as a secondary (non-principal) diagnosis |  | Asthma as either the principal or secondary diagnosis |  |
| --- | --- | --- | --- | --- | --- | --- |
|  | Flu | RSV | Flu | RSV | Flu | RSV |
| <1y | 111.9<br>(91.4,133.1) | 360.1<br>(297.8,421.8) | 5.5<br>(-0.1,11.1) | 158.3<br>(139.4,177) | 6.4<br>(0.1,13) | 209.6<br>(178.2,241.9) |
| 1y | 59<br>(46.5,71.8) | 247.3<br>(207.4,286.8) | 10.1<br>(3.1,17.3) | 143.8<br>(120.2,167.6) | 14.3<br>(4.1,24.8) | 221.4<br>(171.4,270.8) |
| 2y | 36.5<br>(28.7,44.4) | 152.6<br>(126.2,178.3) | 9.3<br>(4.3,14.4) | 96.8<br>(79.4,113.8) | 13.1<br>(6,20.7) | 176.4<br>(137.3,215.7) |
| 3y | 28.6<br>(20.6,36.7) | 94<br>(69,118.9) | 8.1<br>(3.9,12.3) | 58.7<br>(44.4,73.1) | 10.3<br>(5,16) | 107.7<br>(70.9,144.9) |
| 4y | 22.9<br>(17.6,28.4) | 47<br>(29.5,64.4) | 6<br>(2.3,9.8) | 22.9<br>(10.1,35.5) | 9.5<br>(3.5,15.8) | 49.3<br>(12.6,86.7) |
| 5-6y | 21.5<br>(16.8,26.3) | 29.1<br>(13.8,44.3) | 7.4<br>(4.2,10.7) | 16.2<br>(5.3,27.1) | 9.8<br>(5.7,14.4) | 27<br>(-7.2,61.6) |
| 7-11y | 11.2<br>(9.3,13.2) | 10.4<br>(4.2,16.5) | 5.1<br>(3.2,7) | 10.6<br>(4.1,17.2) | 7.3<br>(4.5,10.2) | 20.1<br>(-1.1,41.5) |
| 12-17y | 8.1<br>(7.1,9) | 8<br>(5,11.1) | 2<br>(-1,5.1) | 13.9<br>(3.5,24.2) | 3.4<br>(-1.8,8.5) | 18.6<br>(5,32.3) |

**Table 3.**
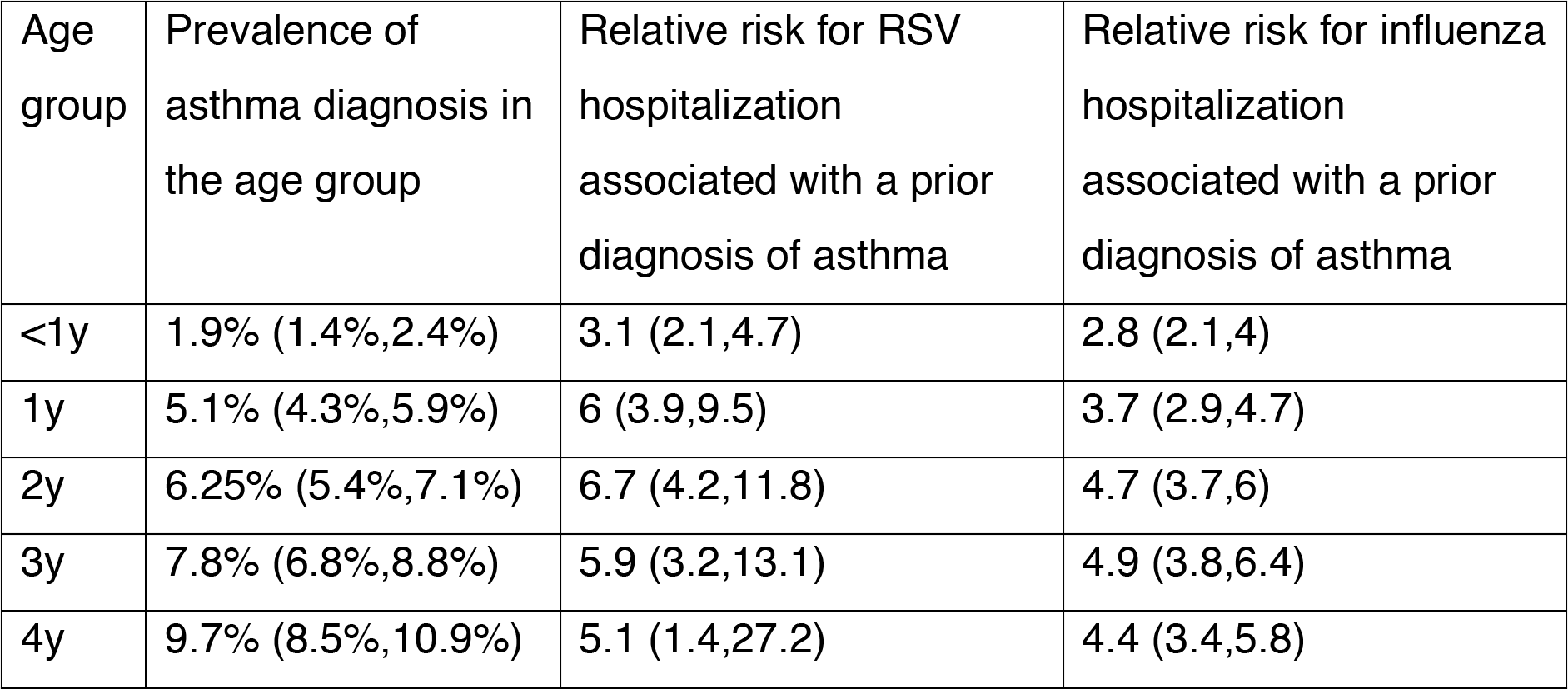
Average annual prevalence of asthma diagnosis in US children aged <5y between 2003-2009, and relative risks for RSV and influenza hospitalization associated with a prior diagnosis of asthma between the 2003-04 through the 2009-10 seasons

Table 3 presents the estimates of the prevalence of asthma by age group between 2003-2009 in young children in the US, and the relative risks for RSV and influenza-associated hospitalization associated with a prior diagnosis of asthma between the 2003-04 through the 2009-10 seasons in the US. Prior diagnosis of asthma is associated with significant risks for both RSV and influenza hospitalization in children aged <5y, with those risks ranging between 3.1(2.1,4.7) (age <1y) to 6.7(4.2,11.8) (age 2y) for RSV hospitalizations, and between 2.8(2.1,4) (age <1y) to 4.9(3.8,6.4) (age 3y) for influenza hospitalizations. Additionally, the estimated relative risks for RSV hospitalization were somewhat higher than the corresponding relative risks for influenza hospitalization.

## Discussion

Granular evaluation of the burden of severe outcomes associated with influenza and RSV in children, including children with underlying health conditions should aid in the design of mitigation efforts. For example, while a number of RSV vaccine candidates are currently in different stages of development [17–19], target groups for RSV vaccination beyond the infant population have not been well characterized. In this paper we apply our previously developed methodology [13,14] to estimate the rates of influenza and RSV-associated hospitalization for various discharge diagnoses by age group in US children between 2003-2010. Estimates of the rates of RSV-associated hospitalization in young children are very high (significantly higher than the corresponding estimates of influenza-associated hospitalization rates) and decline rapidly by year of age, most steeply for the first vs. second year of life. Additionally, our results suggest that having a diagnosis of asthma carries significant risks for both RSV and influenza-associated hospitalization in children. The rates of RSV-associated hospitalization in the youngest children with a prior diagnosis of asthma are exceptionally high, and development of RSV-related prevention strategies in those children is especially critical. Such strategies may include annual vaccination and possibly RSV prophylaxis.

We note that many (presumably most) of the estimated RSV and influenza-associated hospitalizations in children involved complications stemming from bacterial infections, but the course of disease was initially triggered by viral infections. For example, Figure 2 suggests that most hospitalizations for pneumonia and influenza (P&) in young children above the seasonal baseline are explained (well matched) by the patterns of RSV circulation. Influenza circulation also explains some of the P&I hospitalizations above the seasonal baseline in Figure 2, with major influenza epidemics (e.g. the 2003-04 season, and the Fall of 2009) corresponding to spikes in P&I hospitalization rates. We call those hospitalizations RSV and influenza-associated hospitalizations, though the etiology behind those hospitalization outcomes involves additional factors besides viral infections, such as bacterial infections (e.g. secondary infection by *S. pneumoniae*), underlying health conditions, etc.

The connection between RSV infection in early childhood and the subsequent risk of developing recurrent wheezing and asthma has received considerable attention in the literature [29–32]. The contribution of RSV and influenza to hospitalization burden in children with asthma is less well established, partly due to the complexities of asthma diagnoses in very young children [21,22,33]. A number of studies have focused on the contribution of respiratory viruses, particularly the human rhinovirus, to asthma exacerbations [34,35]. At the same time, infections with respiratory viruses, particularly RSV, in children with asthma can result in hospitalizations that have a principal cause other than asthma exacerbation (e.g. Table 2). Quantification of the full hospitalization burden associated with RSV and influenza circulation in children who have a diagnosis of asthma should therefore help inform prevention and treatment strategies in those children. Several studies have suggested that young children with underlying health conditions, including asthma, are disproportionately represented among young children hospitalized with influenza and RSV infections [1,5–7,23,24,36]. Hall et al. [1] and Glezen et al. [5] demonstrated a high frequency of RSV infections among respiratory hospitalizations in young children, as well as the relatively high proportions of asthma or other pulmonary conditions in RSV-infected hospitalized children. A large Danish study demonstrated that previous asthma hospitalization carried ∼5-fold risk for hospitalization with a confirmed RSV infection before the age of 2 years [23]. Our estimates of the relative risks for both RSV and influenza hospitalization associated with a prior diagnosis of asthma are in line with the findings in other studies [1,5–7,23]. While the etiology behind an asthma diagnosis in young children may be subject to uncertainty [21,22], our results support the notion that an asthma diagnosis serves as a marker for the risk of an RSV and influenza-associated hospitalization in young children. These results suggest that vaccination could be particularly beneficial for young children having a diagnosis of asthma, stressing the need for adherence to annual influenza vaccination recommendations for those children and supporting the potential utility of annual RSV vaccination in those children, even if RSV vaccination in the general population of young children is less frequent. Furthermore, rates of RSV-associated hospitalization in the youngest children having a diagnosis of asthma are exceptionally high, suggesting that immunoprophylaxis for those children during RSV epidemics could be considered.

This study has some limitations. Information on prior asthma diagnosis in hospitalized children was unavailable [25]. Instead, we extrapolated the relation between frequencies of a prior diagnosis of asthma vs. the hospital discharge diagnosis of asthma for cases of RSV-associated hospitalization from analogous data on influenza hospitalizations reported to FluSurv-NET surveillance – see Supporting Information. While such extrapolation is imperfect, we note that relatively high proportions of cases of influenza-related hospitalization in different age subgroups of children in FluSurv-NET that have asthma listed on the discharge diagnosis also have a prior diagnosis of asthma, and vice versa, suggesting a strong connection between those two categories of hospitalization. Our estimates of several quantities, such as the risk for influenza hospitalization associated with having a diagnosis of asthma rely on how accurately the FluSurv-NET surveillance data represent certain quantities (such as the proportion of hospitalizations that have asthma as the principal discharge diagnosis) for all influenza-related hospitalizations in children (Supporting Information). Another potential issue is the accuracy of the proposed regression framework for the hospitalization time-series [13,14]. Spikes in hospitalization rates during certain influenza seasons correspond visually to major influenza epidemics (as suggested by the incidence proxies that we utilize), providing additional support for the validity of our inference method for influenza-associated hospitalization rates. While RSV circulation is more periodic than influenza circulation, there is notable aperiodicity in the RSV circulation (Figure S1 in the Supporting Information); use of a flexible model for the baseline rates of hospitalization not associated with influenza or RSV helps separate the baseline rates from the rates of hospitalization associated with influenza and RSV and results in significant improvement in the model fits compared to the use of a trigonometric model for the baseline rates (Supporting Information for [14,13]). We used rates of hospitalizations in infants with RSV in the principal diagnosis as a proxy for RSV incidence, while presence of RSV in the principal diagnosis need not be supported by a laboratory test. We excluded certain states with unusual patterns of hospitalization rates in infants with RSV in the principal diagnosis from the analyses (see Supporting Information) in an effort to minimize the impact of variable quality of the proxy. We note that our models produce good, temporally consistent fits to the data (R-squared statistic above 0.98 for all models, Supporting Information), especially for the younger children (e.g. Figures 1-3). Additionally, the confidence bounds for the contribution of RSV to many of the hospitalization categories considered are reasonably tight (Tables 1 and 2), and our estimates for the rates of RSV-associated hospitalization are generally consistent with the ones in the literature [8,9]. Finally, the estimated weekly baseline rates of hospitalization not associated with RSV and influenza exhibit Fall and Spring peaks (Figures 1-3) that are suggestive of the contribution of rhinoviruses. Further related work is needed to examine various aspects of our inference methodology, including the effect of the circulation of other respiratory viruses.

We believe that despite those limitations, our work provides granular estimates of the rates of hospitalization associated with influenza virus and RSV infections in children, suggesting very high rates of RSV-associated hospitalizations in young children. Such work, in conjunction with efforts to understand the role of different age groups in propagating RSV epidemics [37] may aid the evaluation of the impact of RSV vaccination on disease rates in different population groups, including children, and inform RSV vaccination strategies. Additionally, our results demonstrate risks for both RSV and influenza hospitalization associated with a prior diagnosis of asthma in young children, with the rates of RSV-related hospitalization in the youngest children having a diagnosis of asthma (particularly those aged <2y) being especially high. These results may help inform prevention efforts such as vaccination strategies for young children having a diagnosis of asthma during RSV and influenza epidemics, and possibly RSV-related prophylaxis for the youngest children having a diagnosis of asthma.

## Acknowledgement

We would like to thank the HCUP Partner states that voluntarily provide their data to the project (https://www.hcup-us.ahrq.gov/db/state/siddbdocumentation.jsp). We are also grateful to Dr. Lara Akinbami of the US CDC who provided data on the prevalence of asthma by year of age in the US children during the study period. The findings and conclusions in this report are those of the authors and do not necessarily represent the official position of the Centers for Disease Control and Prevention.

## Supporting Information for: “Hospitalizations associated with respiratory syncytial virus RSV) and influenza in children, including children having a diagnosis of asthma”

### Section S1: Hospitalization categories used in the analyses

Table S1 lists the six hospitalization categories used in the analyses

**Table S1:** Hospitalization categories used in the analyses

| No. | Category | ICD-9 codes |
| --- | --- | --- |
| 1 | Respiratory cause excluding asthma in the principal discharge diagnosis | 460-519, but not 493 in principal discharge diagnosis |
| 2 | Respiratory cause present anywhere in the discharge diagnoses excluding asthma in the principal diagnosis | 460-519 anywhere on discharge diagnosis, but not 493 in principal discharge diagnosis |
| 3 | Respiratory cause present anywhere in the discharge diagnosis | 460-519 anywhere on discharge diagnosis |
| 4 | Pneumonia and influenza in the principal discharge diagnosis | 480-488 in the principal discharge diagnosis |
| 5 | Asthma as a secondary (non-principal) discharge diagnosis | 493 as a non-principal discharge diagnosis |
| 6 | Asthma present anywhere in the discharge diagnosis | 493 anywhere in the discharge diagnosis |

### Section S2: Incidence proxies for the major influenza (sub)types, and RSV

As the rates of influenza and RSV infection are difficult to estimate directly, we derive *proxies* for the incidence of RSV and the major influenza (sub)types (A/H3N2, A/H1N1, and influenza B) that are expected to be proportional to the population incidence of those viruses.

For the RSV incidence proxy, we utilize data on hospitalizations with the principal diagnosis of RSV (ICD-9 codes 466.11, 480.1, and 079.6) in infants aged <1y reported to the State Inpatient Databases of the Healthcare Cost and Utilization Project (HCUP) [1]. We use hospitalization rates in infants because there is an apparent upward trend in annual rates of hospitalization with the principal diagnosis of RSV in all age groups above two years of age in the US, likely due to changes in testing/diagnostic practices, but no apparent trend in the rates of hospitalization with the principal diagnosis of RSV in infants. Additionally, the vast majority of RSV hospitalizations in infants are for RSV bronchiolitis (ICD-9 code 466.11). We believe that rates of RSV bronchiolitis in infants were proportional to the rates of RSV infection in infants, at least up to mid 2000s or so, because of evidence in the literature (e.g. [2]) that the vast majority of bronchiolitis diagnoses were supported by laboratory tests during that time. We note that for this study, rates of RSV bronchiolitis in infants were not available. Later on, testing practices for viral infections following a bronchiolitis diagnosis were changing over time, and laboratory testing was no longer recommended in the US by 2014 [3]. Moreover, location-specific differences in the practices for assigning the principal discharge diagnosis of RSV in infants could have existed even before that. Indeed, annual rates of states than the corresponding rates in most states, while certain other states exhibited apparent temporal trends in the rates of hospitalization with RSV in the principal diagnosis in infants. In an effort to reduce potential biases in the relation between the rates of RSV hospitalization diagnoses in infants and RSV population incidence rates, the following states were excluded from our analyses: (i) states with an apparent temporal trend in the rates of hospitalization in infants with RSV in the principal diagnosis (three states); (ii) states that did not report RSV hospitalizations in infants during certain seasons (one state); (iii) states for which annual rates of hospitalization in infants with RSV in the principal diagnosis were noticeably higher than in most states (seven states). Overall, among the 35 states that reported hospitalization data to HCUP between 2003-2010, 24 were included in our analyses. Those states are “AR” “CA” “CO” “CT” “GA” “HI” “IA” “IL” “IN” “MD” “MN” “NC” “NE” “NJ” “NV” “NY” “OH” “OR” “TN” “TX” “VA” “VT” “WA” “WI”. We consider the weekly incidence proxy for RSV for the collection of states included in our analyses to be the combined rate of hospitalization in those states with the principal diagnosis of RSV per 100,000 children aged <1y during that week.

The incidence proxies for the major influenza (sub)types are calculated from the US CDC surveillance data [4]. For each US state, we combined weekly data for that state on medical consultations in the US CDC Outpatient Illness Surveillance Network (ILINet), including those that were for influenza-like illness (ILI) with data on testing of respiratory specimens in the US Virologic Surveillance laboratories to define weekly proxies for the incidence of each major influenza (sub)type in the given state. Specifically,

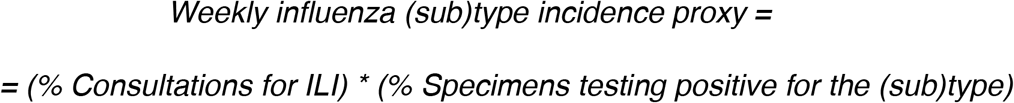

The (sub)type-specific incidence proxy for the collection of states used in our analyses was defined as the sum of state-specific incidence proxies for the (sub)type weighted by the weekly state populations, with state populations during various weeks estimated through linear interpolation (in time) for the yearly, July 1st state population estimates [5].

Figure S1 shows the incidence proxies for influenza A/H3N2, A/H1N1, and B, as well as for RSV. The incidence proxy for RSV was more periodic than the incidence proxies for the different influenza (sub)types, with the highest cumulative incidence for the RSV incidence proxy attained during the 2003-04 season.

**Figure S1:**
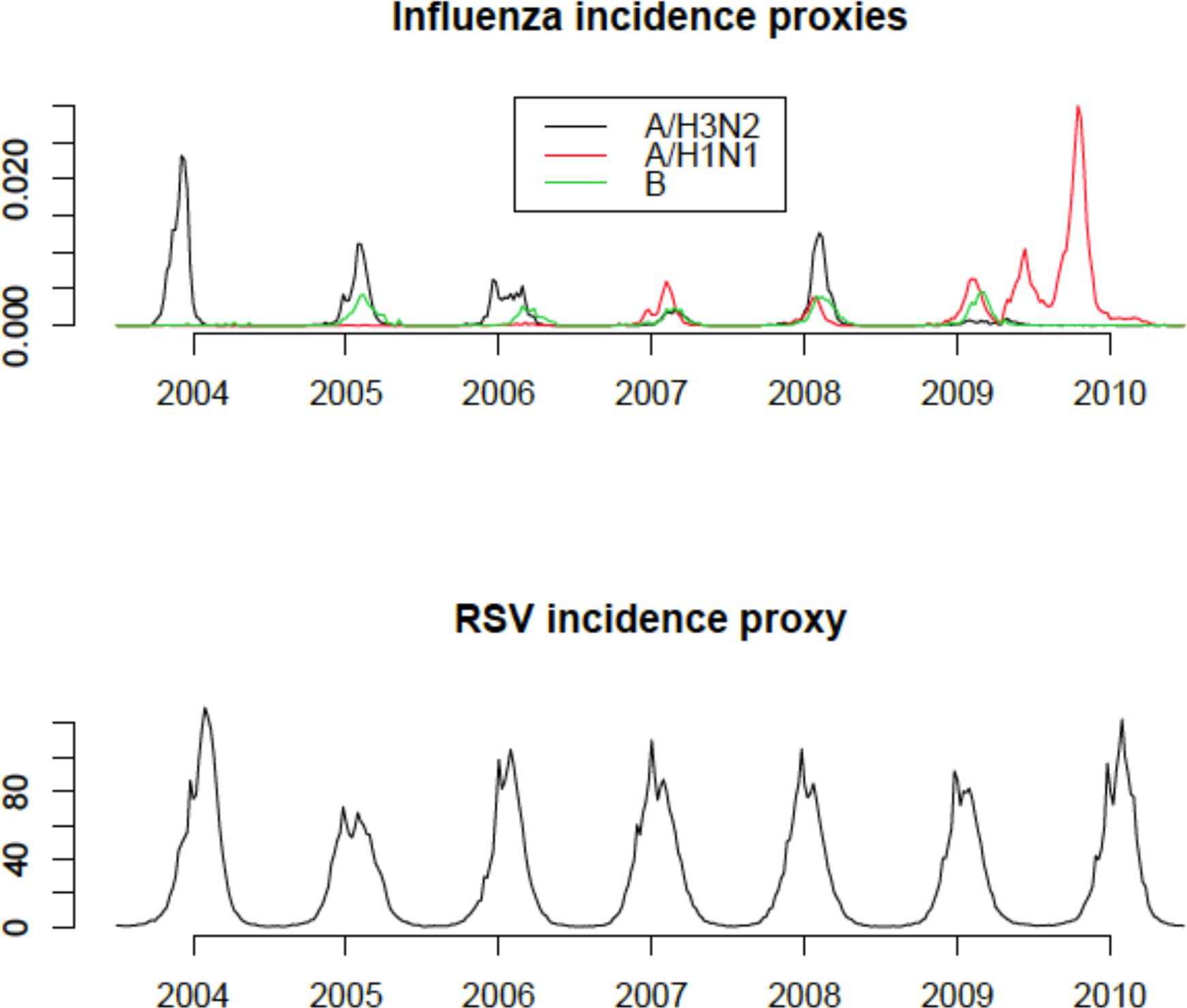
Incidence proxies for influenza A/H3N2, A/H1N2, and B (top panel), and for RSV (bottom panel).

The relationship between an influenza incidence proxy and the rate of associated hospitalization in a given age group may change in time due to a variety of factors [6]. One of those factors is change in the age distribution for different influenza-associated outcomes due to changes in the circulating influenza strain and in population immunity. In particular, the 2003-04 influenza season in the US was driven by a novel (Fujian) A/H3N2 strain, with the A/H3N2 incidence proxy for that season being exceptionally large (Figure S1). The relation (ratio) between the A/H3N2 incidence proxy (Figure S1) and A/H3N2 influenza incidence in young children for the 2003-04 season could be different compared to other seasons. This notion is supported by the fact that school-age children are overrepresented in influenza incidence (and surveillance data) during the circulation of novel influenza strains compared to other seasons [7,8]; moreover, higher rates of excess respiratory ED visits in school-age compared to pre-school children were estimated during the 2003-04 season compared to the 2004-05 season in NYC [9]. Similarly, the emergence of the pandemic A/H1N1 influenza strain in 2009 resulted in exceptionally high rates of pediatric influenza infections [7], and the ratios between the rates of A/H1N1-associated hospitalizations in different subgroups of children and the A/H1N1 incidence proxy are potentially different for the pandemic strain compared to the distantly-related seasonal A/H1N1 strain in preceding seasons. To address those considerations, we split the A/H3N2 incidence proxy into two, namely H31 equaling the A/H3N2 proxy for the 2003-04 season, with H31 being zero afterwards, and H32, being zero for the 2003-04 season, and equaling the A/H3N2 incidence proxy afterwards. Similarly, we split the A/H1N1 incidence proxy into two, reflecting A/H1N1 incidence before and during the 2009 pandemic.

### Section S3: Model fits

In this section we show model fits for hospitalizations with a respiratory cause present anywhere in the discharge diagnosis (excluding the principal diagnosis of asthma), pneumonia and influenza in the principal diagnosis, asthma as a secondary diagnosis, and asthma present anywhere in the discharge diagnosis in US children from the 2003-04 through the 2009-10 seasons. The plots suggest that the model fits were generally good and temporally consistent (all R-squared statistics being above 0.98, Table S2), particularly for the younger children. Model fits for respiratory hospitalizations were generally better compared to hospitalizations with asthma present anywhere in the discharge diagnosis (Table S2). Additionally, there are double seasonal peaks (Fall and Spring) for the baseline rates of hospitalization not associated with RSV or influenza for a number of hospitalization categories and age groups (particularly non-infants). Those double peaks may partly reflect the contribution of rhinoviruses to the corresponding hospitalization burden [10–12].

**Table S2:** R-squared statistic for the model fits for different age groups/categories of hospitalization

|  | <1y | 1y | 2y | 3y | 4y | 5-6y | 7-11y | 12-17y |
| --- | --- | --- | --- | --- | --- | --- | --- | --- |
| Respiratory cause (excluding asthma) in the principal diagnosis | 0.999 | 0.994 | 0.992 | 0.99 | 0.989 | 0.987 | 0.989 | 0.991 |
| Respiratory present anywhere in the discharge diagnosis excluding asthma in the principal diagnosis | 0.999 | 0.995 | 0.994 | 0.992 | 0.992 | 0.992 | 0.993 | 0.993 |
| Respiratory cause present anywhere in the discharge diagnosis | 0.999 | 0.996 | 0.994 | 0.992 | 0.991 | 0.989 | 0.991 | 0.993 |
| Pneumonia and influenza in the principal discharge diagnosis | 0.992 | 0.992 | 0.989 | 0.985 | 0.984 | 0.981 | 0.985 | 0.985 |
| Asthma as a secondary diagnosis | 0.994 | 0.993 | 0.991 | 0.987 | 0.986 | 0.986 | 0.991 | 0.989 |
| Asthma present anywhere in the discharge diagnosis | 0.994 | 0.993 | 0.99 | 0.987 | 0.983 | 0.98 | 0.985 | 0.99 |

#### Section S3.1: Hospitalizations with a respiratory cause in either the principal or secondary diagnosis (excluding the principal diagnosis of asthma)

**Figure S2:**
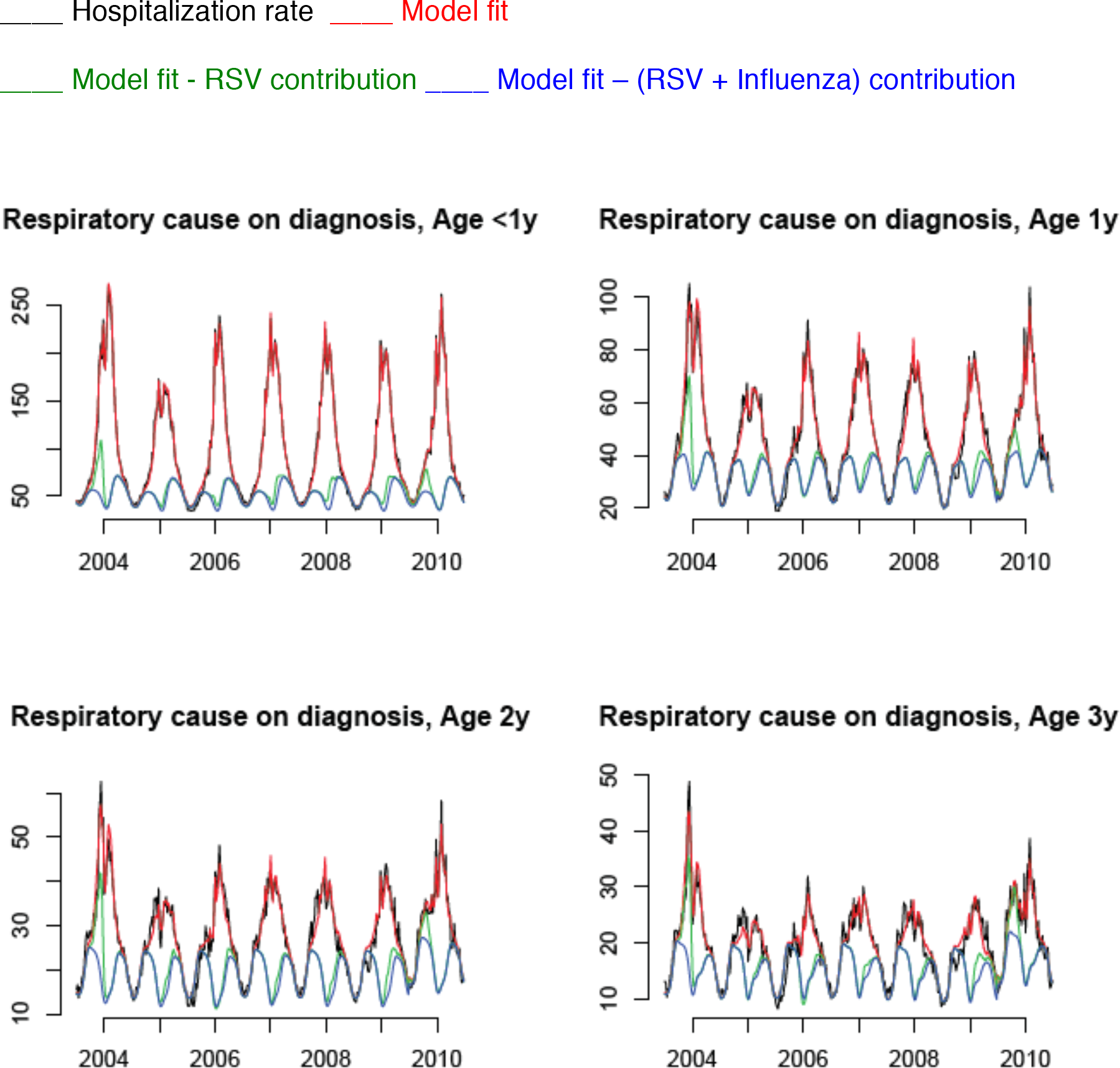
Weekly hospitalization rates (per 100,000) with a respiratory cause present anywhere in the discharge diagnosis for children aged <4y, 2003-04 through the 2009-10 seasons (black), model fits (red), and contributions of RSV (red curve minus green curve) and influenza (green curve minus blue curve).

**Figure S3:**
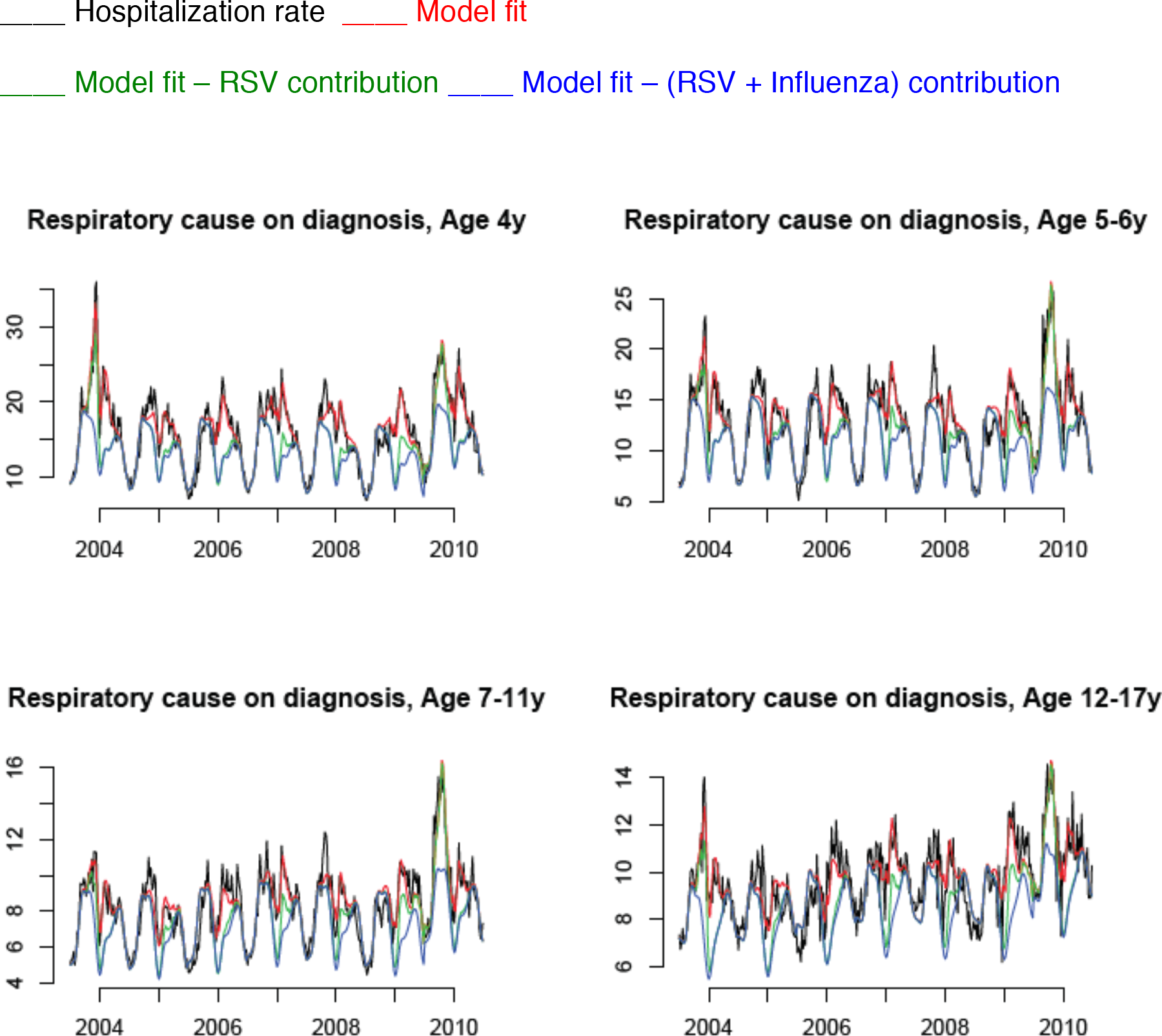
Weekly hospitalization rates (per 100,000) with a respiratory cause present anywhere in the discharge diagnosis for children aged 4-17y, 2003-04 through the 2009-10 seasons (black), model fits (red), and contributions of RSV (red curve minus green curve) and influenza (green curve minus blue curve).

#### Section S3.2: Hospitalizations with pneumonia and influenza in the principal diagnosis

**Figure S4:**
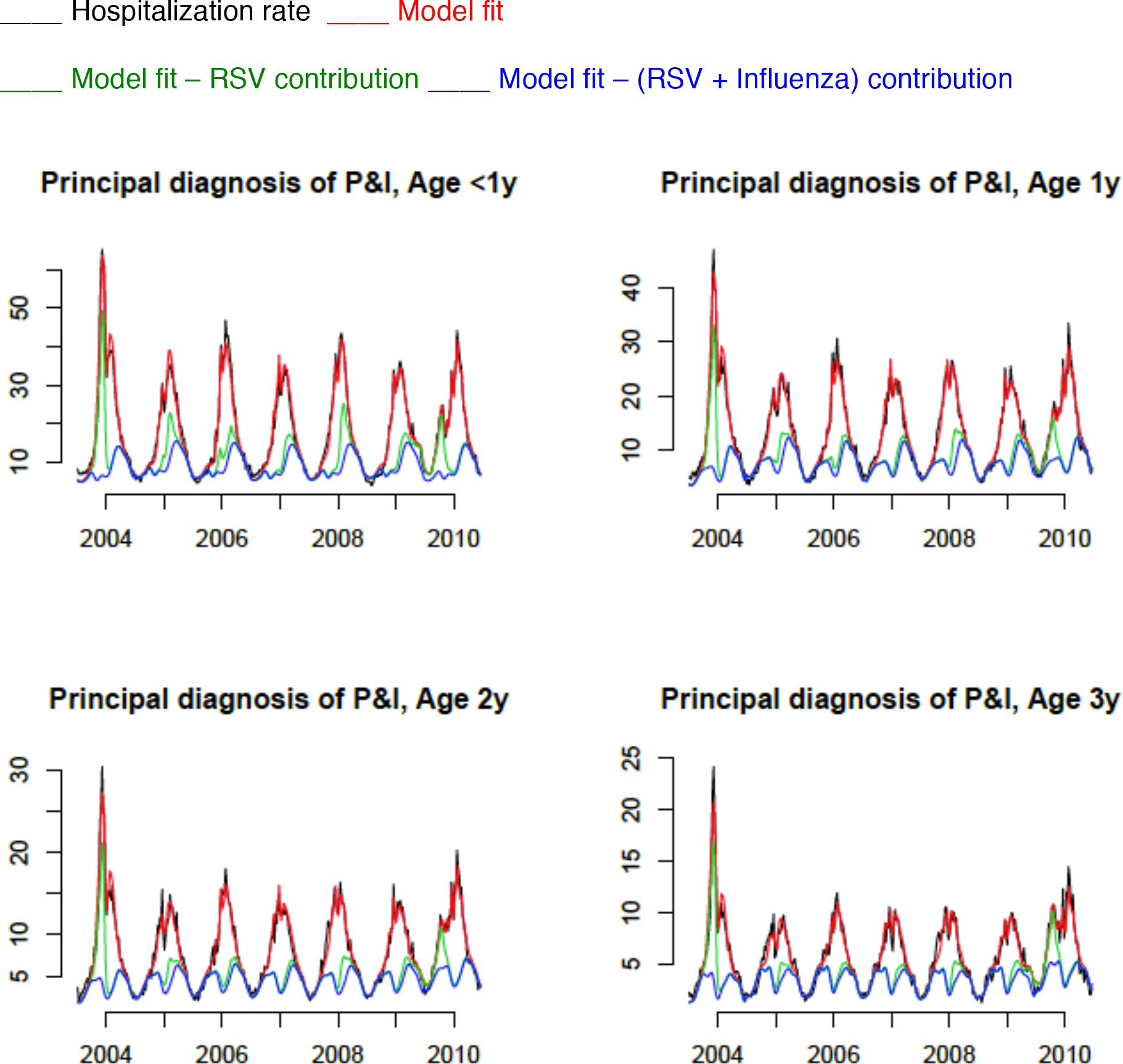
Weekly hospitalization rates (per 100,000) with pneumonia and influenza in the principal discharge diagnosis for children aged <4y, 2003-04 through the 2009-10 seasons (black), model fits (red), and contributions of RSV (red curve minus green curve) and influenza (green curve minus blue curve).

**Figure S5:**
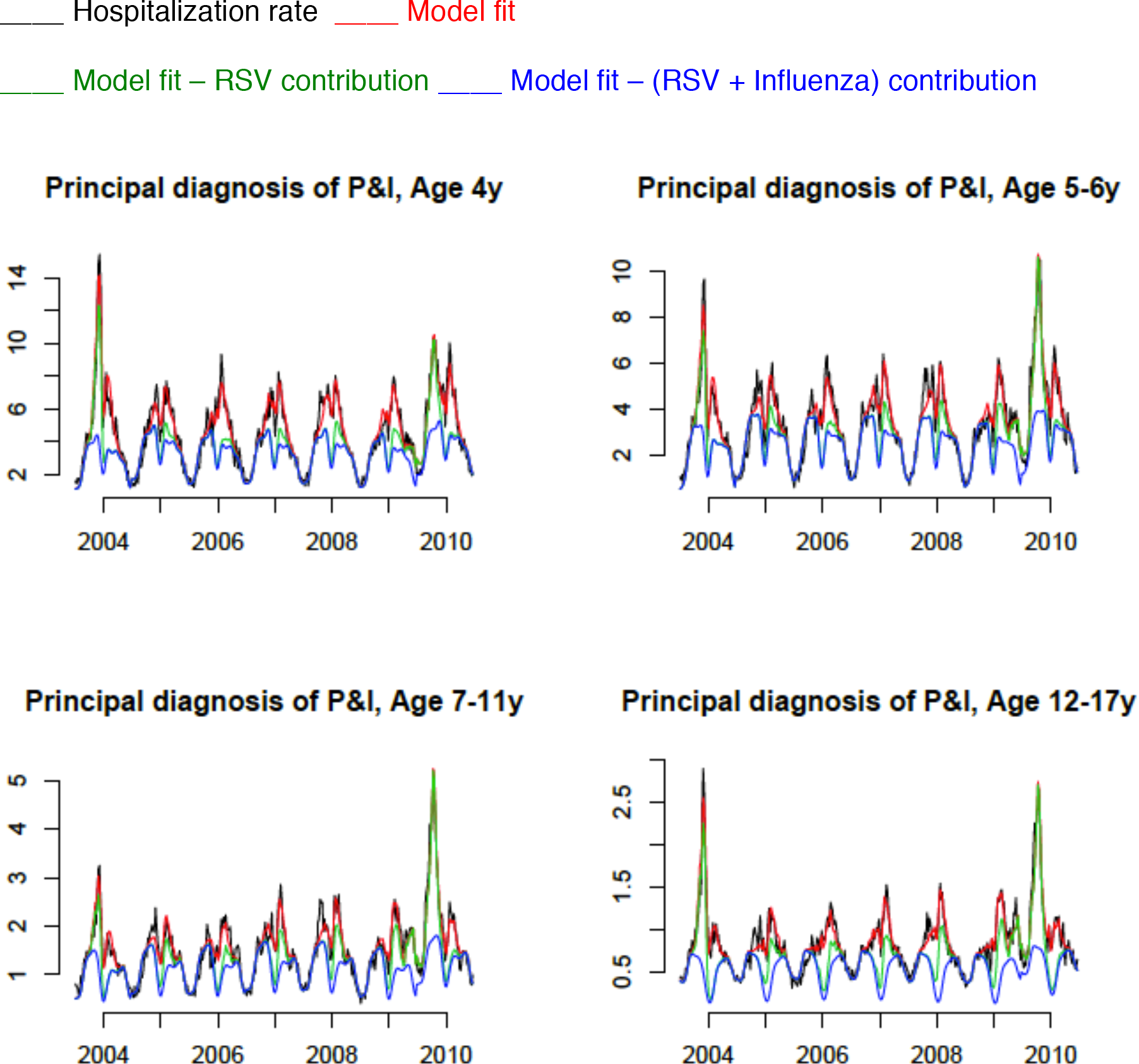
Weekly hospitalization rates (per 100,000) with pneumonia and influenza in the principal discharge diagnosis for children aged 4-17y, 2003-04 through the 2009-10 seasons (black), model fits (red), and contributions of RSV (red curve minus green curve) and influenza (green curve minus blue curve)

#### Section S3.3: Hospitalizations with asthma as a secondary diagnosis

**Figure S6:**
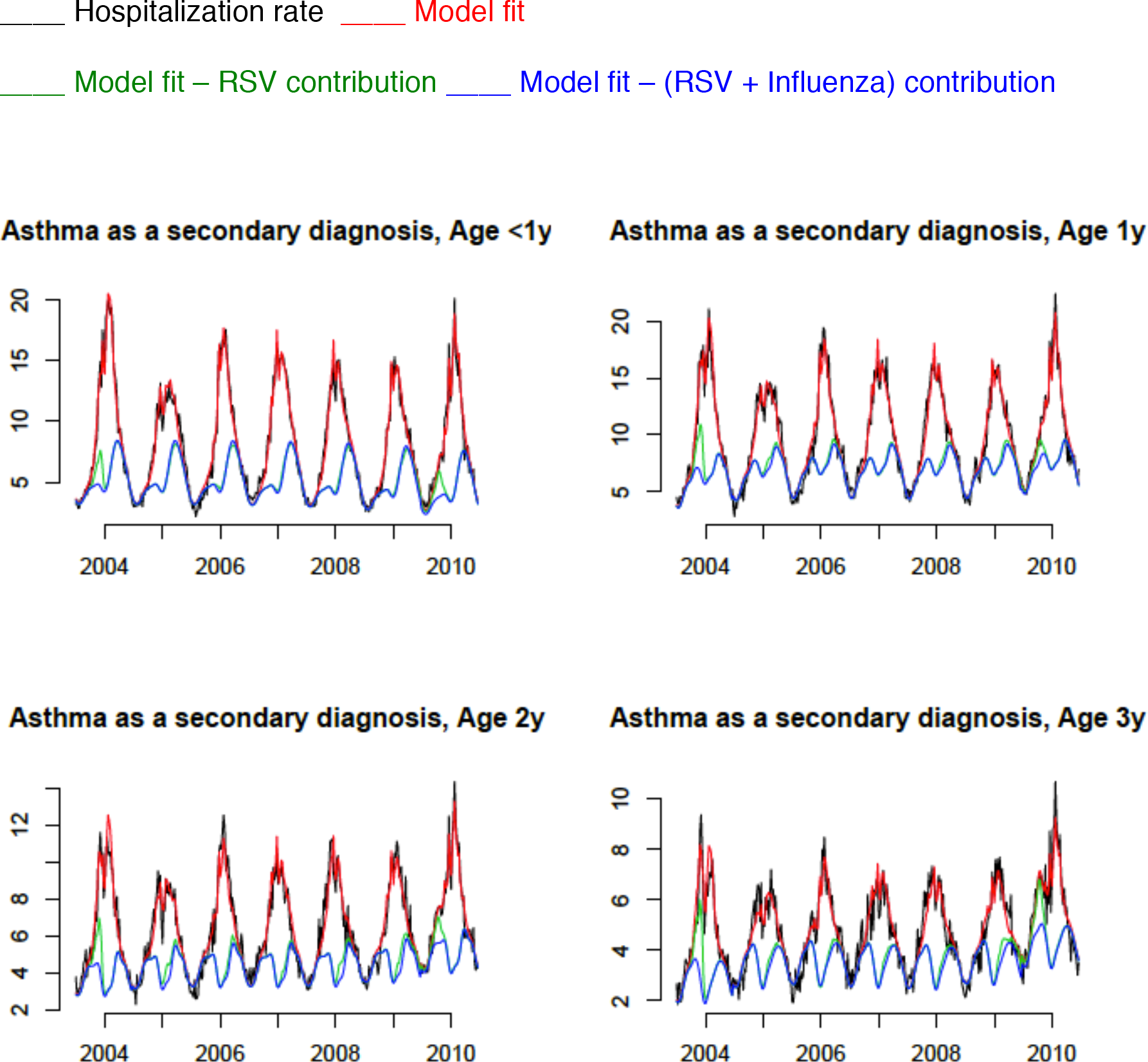
Weekly hospitalization rates (per 100,000) with asthma as a secondary discharge diagnosis for children aged <4y, 2003-04 through the 2009-10 seasons (black), model fits (red), and contributions of RSV (red curve minus green curve) and influenza (green curve minus blue curve).

**Figure S7:**
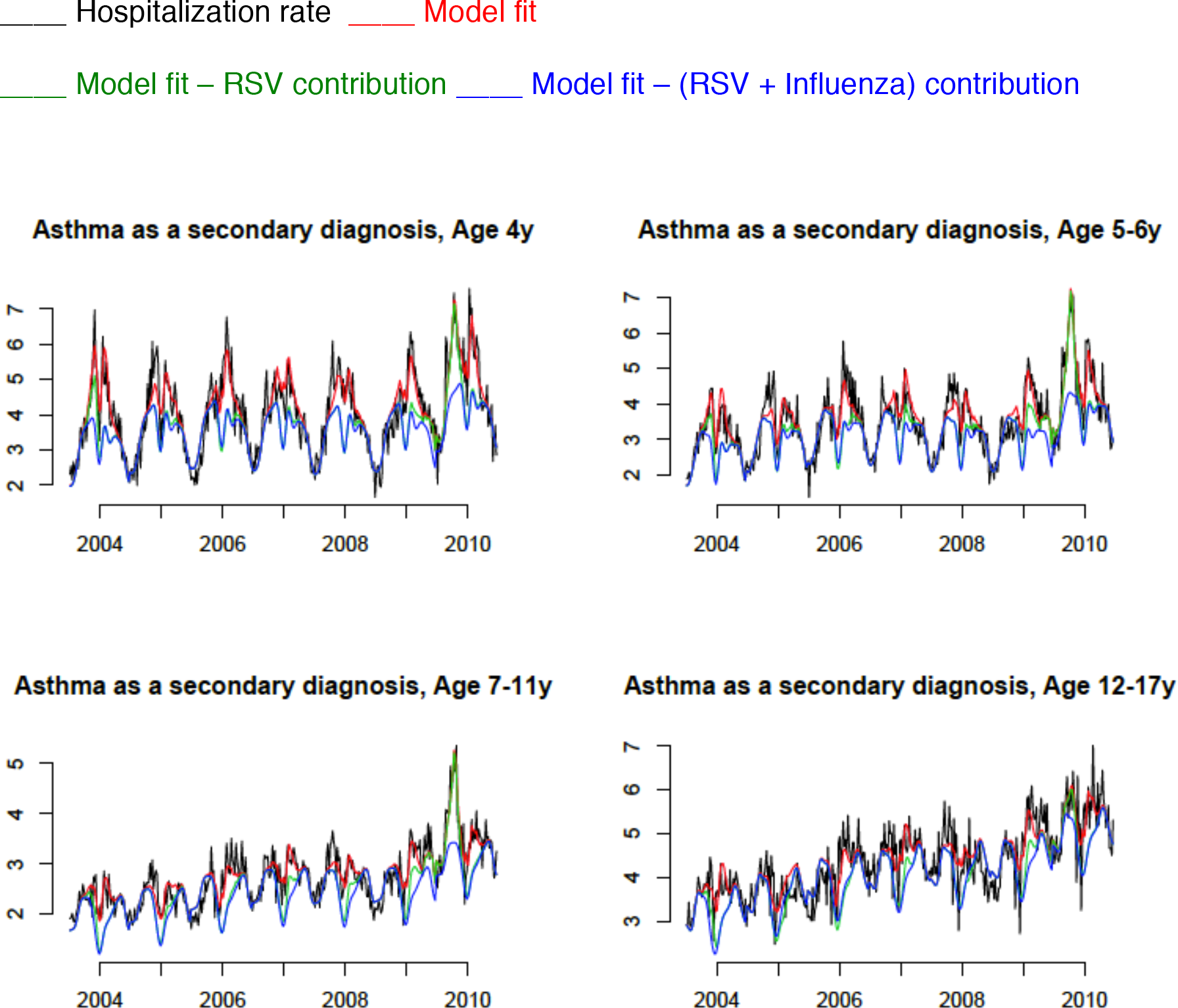
Weekly hospitalization rates (per 100,000) with asthma as a secondary discharge diagnosis for children aged 4-17y, 2003-04 through the 2009-10 seasons (black), model fits (red), and contributions of RSV (red curve minus green curve) and influenza (green curve minus blue curve).

#### Section S3.4: Hospitalizations with asthma present anywhere in the discharge diagnosis

**Figure S8:**
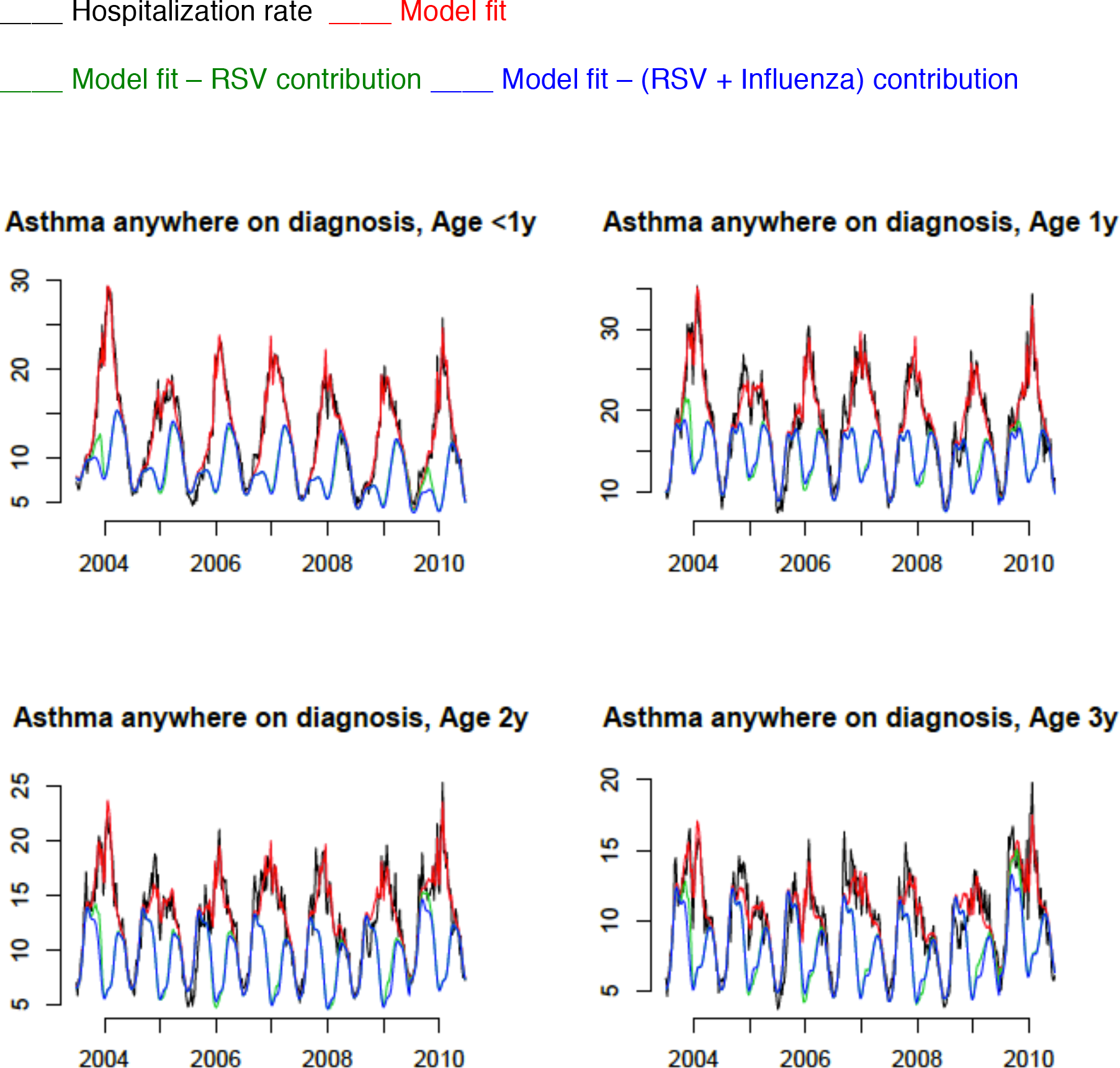
Weekly hospitalization rates (per 100,000) with asthma present anywhere in the discharge diagnosis for children aged <4y, 2003-04 through the 2009-10 seasons (black), model fits (red), and contributions of RSV (red curve minus green curve) and influenza (green curve minus blue curve)

**Figure S9:**
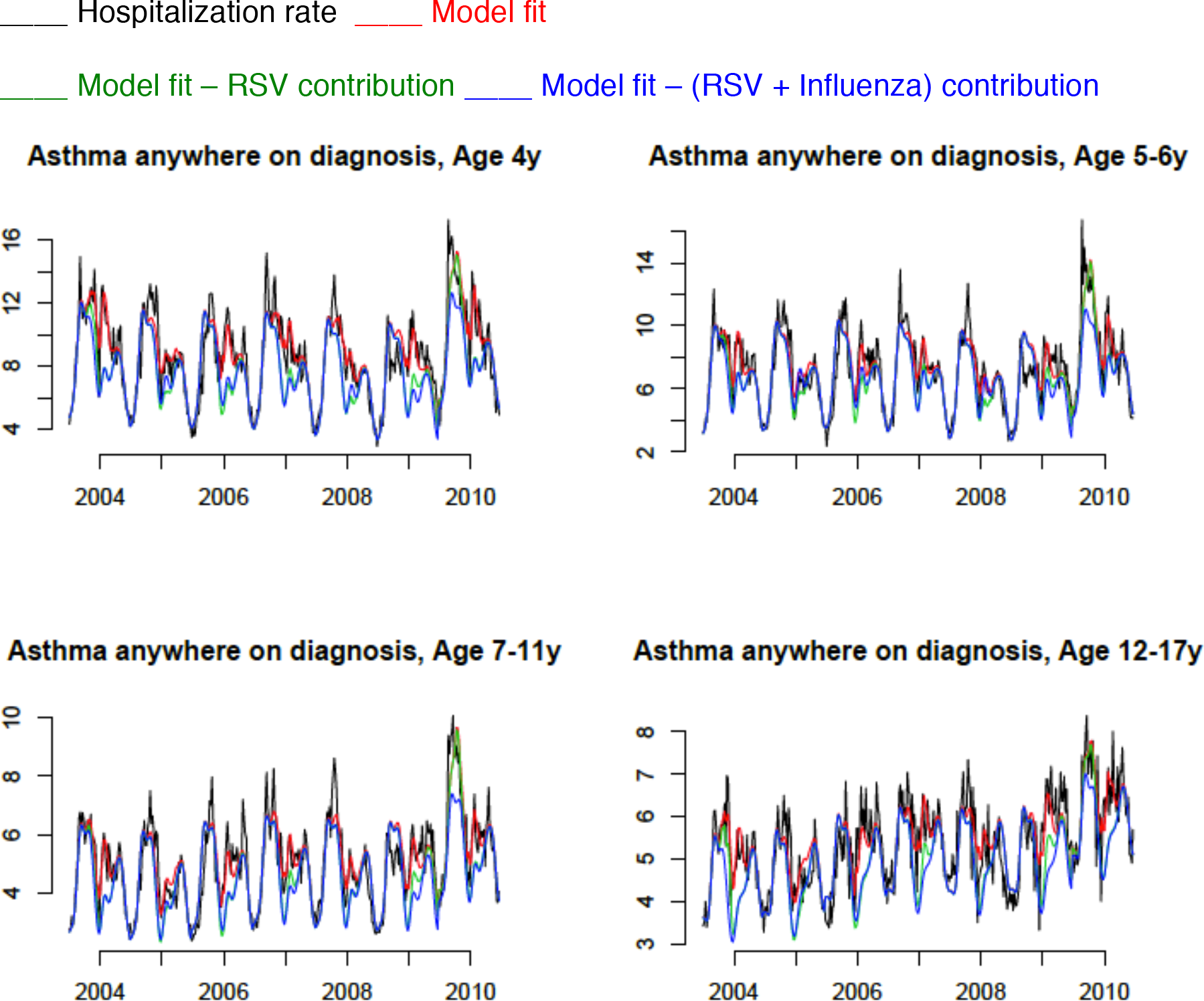
Weekly hospitalization rates (per 100,000) with asthma present anywhere in the discharge diagnosis for children aged 4-17y, 2003-04 through the 2009-10 seasons (black), model fits (red), and contributions of RSV (red curve minus green curve) and influenza (green curve minus blue curve).

### Section S4: Rates of influenza-associated hospitalization with a respiratory cause present in the discharge diagnosis, as well as with asthma present in the discharge diagnosis

The rates of hospitalization in category 3 in the Methods section (respiratory cause present anywhere in discharge diagnosis) are the sum of hospitalization rates in category 2 (respiratory cause present anywhere in the discharge diagnosis excluding the principal discharge diagnosis of asthma) and rates of hospitalization with asthma as the principal discharge diagnosis. The weekly rates of hospitalization with asthma as the principal discharge diagnosis are not periodic year-to-year. Moreover, this aperiodicity is correlated with the aperiodicity in the incidence proxies for the major influenza subtypes, and this biases the estimates of the contribution of influenza to the rates of hospitalization with a respiratory cause present anywhere in the discharge diagnosis (category 3). In this section, we present a different estimation of the contribution of influenza to the rates of hospitalization with a respiratory cause present anywhere in the discharge diagnosis (category 3). This inference utilizes data on FluSurv-NET influenza-related hospitalizations in children [13,14]. Those surveillance data cover about 9% of the US population and represent laboratory-confirmed influenza hospitalizations. The estimation for category 3 is performed under the assumption that the proportion of influenza-associated hospitalizations with asthma as the principal discharge diagnosis among all influenza-associated hospitalizations in category 3 equals the proportion of influenza-related hospitalizations in FluSurv-NET [13,14] between 2003-04 through 2009-10 seasons that have asthma as the principal discharge diagnosis. Specifically, to estimate the rates of influenza-associated hospitalization in category 3, we

A. Estimate of the age-specific rate of influenza-associated hospitalization with a respiratory cause present anywhere in the discharge diagnosis excluding the principal diagnosis of asthma (category 2 in the Methods)
B. Estimate the age-specific proportion of influenza-related hospitalizations in the FluSurv-NET surveillance database [13,14] between 2003-04 through 2009-10 seasons that have asthma as the principal discharge diagnosis
C. Divide the estimate in A) by (1 – minus the estimate in B))

Rates of influenza-associated hospitalization with asthma present anywhere in the discharge diagnosis (category 6 in the Methods section) are estimated analogously using the estimates of the rates of influenza-associated hospitalization with asthma as a secondary (non-principal) discharge diagnosis (Table 2 in the main text), and the proportion of FluSurv-NET hospitalizations [13,14] with asthma present anywhere in the discharge diagnosis that have asthma as the principal discharge diagnosis.

### Section S5: Relative risks for RSV and influenza hospitalization associated with a prior diagnosis of asthma

For a given age group, let *A* be the number of children with a prior diagnosis of asthma; *B* be the number of children without a prior diagnosis of asthma; *HA*(*flu*) be the number influenza-related hospitalizations in children with a prior diagnosis of asthma; and *HB*(*flu*) be the number of influenza-related hospitalizations associated in children without a prior diagnosis of asthma. The relative risk (risk ratio) for influenza-related hospitalization associated with a prior diagnosis of asthma in a given age group is then

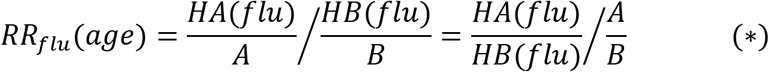

The quantity 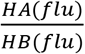 is estimated as 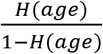, where *H*(*age*) is the (age-specific) proportion of cases of influenza-related hospitalization in the FluSurv-NET surveillance database [13,14] between the 2003-04 through the 2009-10 seasons that have a previous history of asthma. The quantity 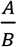 is estimated as 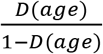, where *D*(*age*) is the age-specific population prevalence of an asthma diagnosis [15]. Data on the prevalence of asthma diagnosis by year of age are described in [15], with the corresponding annual averages between 2003-2009 provided to us by Dr. L. Akinbami of the US CDC. With those estimates, equation (^*^) becomes eq. 3 in the main text.

Estimation of the age-specific risk of RSV hospitalization associated with a prior diagnosis of asthma is performed analogously, namely

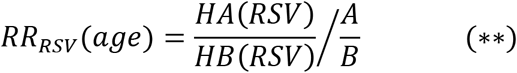

To estimate the quantity *HA*(*RSV*), the (age-specific) number of RSV-related hospitalizations in children having a prior diagnosis of asthma, let *SA*(*RSV*) be the (age specific) number of RSV-related hospitalizations with asthma as a secondary discharge diagnosis. The quantity *SA*(*RSV*) is estimated in Table 2 in the main text. Additionally, let *HA*(*RSV*)&*SA*(*RSV*) be the (age-specific) number of cases of RSV-related hospitalization where the child had both: i) a prior diagnosis of asthma and ii) asthma listed as a secondary discharge diagnosis. Then

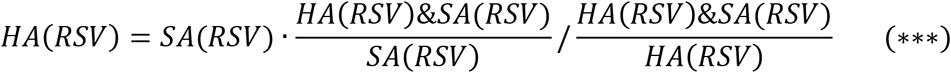

The quantit 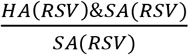 is the (age-specific) probability of having a prior diagnosis of asthma for a case of RSV-related hospitalization with asthma as a secondary discharge diagnosis; the quantity 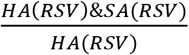 is the (age-specific) probability of having asthma as a secondary discharge diagnosis for a case of RSV-related hospitalization with a prior diagnosis of asthma. While we don’t have data to estimate those conditional probabilities for RSV-related hospitalizations, we adopt the estimates of the corresponding conditional probabilities for influenza-related hospitalizations recorded in the FluSurv-NET data [13,14]. We note that we use secondary diagnosis of asthma rather than any discharge diagnosis of asthma in eq. (^***^) because the probability of symptoms of asthma exacerbation/principal discharge diagnosis of asthma for children without a prior diagnosis of asthma may be different (possibly higher) for an RSV-related hospitalization compared to an influenza-related hospitalization. The estimates of the corresponding conditional probabilities in eq. (***) for an influenza-related hospitalization are presented in Table S3 below.

**Table S3:** Age-specific proportions of children with a history of asthma among hospitalized children with a secondary discharge diagnosis of asthma, as well as proportions of children with a secondary discharge diagnosis of asthma among hospitalized children with a history of asthma.

| Age | Asthma history among cases of hospitalization with asthma as a secondary diagnosis | Secondary asthma diagnosis among case of hospitalization with asthma history |
| --- | --- | --- |
| <1y | 37.8% (29.6%,46.5%) | 45.1% (35.8%,54.8%) |
| 1y | 59.1% (50.2%,67.6%) | 48.8% (40.8%,56.8%) |
| 2y | 65% (55%,74.2%) | 51.5% (42.6%,62.4%) |
| 3y | 71.4% (61.4%,80.1%) | 59.8% (50.4%,68.8%) |
| 4y | 73% (60.3%,83.4%) | 42.6% (33.1%,52.5%) |

### Section S6: Data collection instruments

#### Hospitalization data

Data on hospitalizations for different discharge diagnoses (Table S1) between calendar week 27 of 2003, and calendar week 26 of 2010 are stratified by state/week/age group (<1y,1y,2y,3y,4y,5-6y,7-11y,12-17y). Those data can be requested from the State Inpatient Databases of the Healthcare Cost and Utilization Project (HCUP), maintained by the Agency for Healthcare Research and Quality (AHRQ) through an active collaboration [1].

*Influenza surveillance data:* For the influenza incidence proxies, we utilized the US Centers for Disease Control and Prevention (CDC) influenza surveillance data between calendar week 26 of 2003, and calendar week 26 of 2010 [4]. Two data streams were used in the inference: (i) weekly state-specific percent of medical consultations in the CDC Outpatient Illness Surveillance Network (ILINet) that were for influenza-like illness (ILI);(ii) the state-specific percent of respiratory specimens in the US Virologic Surveillance laboratories that tested positive for each of the major influenza (sub)types (A/H3N2, A/H1N1, and influenza B).

*RSV hospitalization data:* Counts of hospitalization among infants (aged <1y) with RSV (ICD-9 codes 466.11, 480.1, and 079.6 present in the principal discharge diagnosis) between calendar week 27 of 2003, and calendar week 26 of 2010 can be requested from the State Inpatient Databases of the Healthcare Cost and Utilization Project (HCUP), maintained by the Agency for Healthcare Research and Quality (AHRQ) through an active collaboration [1]. We’ve obtained those data stratified by state/week.

Population data: Yearly population counts in different age groups (<1y,1y,2y,3y,4y,5-6y,7-11y,12-17y) between 2003 and 2010 can be obtained from the CDC Wonder website [16]. Those data are stratified by state.

